# An integrative machine learning approach for prediction of toxicity-related drug safety

**DOI:** 10.1101/455667

**Authors:** Artem Lysenko, Alok Sharma, Keith A Boroevich, Tatsuhiko Tsunoda

## Abstract

Recent trends in drug development have been marked by diminishing returns of escalating costs and falling rate of new drug approval. Unacceptable drug toxicity is a substantial cause of drug failure during clinical trials as well as the leading cause of drug withdraws after release to market. Computational methods capable of predicting these failures can reduce waste of resources and time devoted to the investigation of compounds that ultimately fail. We propose an original machine learning method that leverages identity of drug targets and off-targets, functional impact score computed from Gene Ontology annotations, and biological network data to predict drug toxicity. We demonstrate that our method (TargeTox) can distinguish potentially idiosyncratically toxic drugs from safe drugs and is also suitable for speculative evaluation of different target sets to support the design of optimal low-toxicity combinations.

**Summary blurb:** Prediction of toxicity-related drug clinical trial failures, withdrawals from market and idiosyncratic toxicity risk by combining biological network analysis with machine learning.

## Introduction

Last decade has seen an escalation of drug development costs and at the same time, the rate at which new successful drugs are released has actually decreased [1]. One striking example of this trend was put forward by [2], who observed that during the period of 2004-2014, both the funding and number of drug candidates trialed in the US increased substantially, but the number of new drugs approved declined by over 25% compared to the previous decade. Unacceptably high toxicity is a major contributing cause of drug failure and accounts for about one-fifth of clinical trial failures [3] and two-thirds of worldwide post-launch withdrawals [4]. One strategy to reduce these costs and improve the efficiency of the drug development is to augment laboratory and clinical testing with computational analysis [5] and the development of accurate methods to predict toxicity is pivotal to this goal [6].

Earlier methods for computational pre-screening focused primarily on chemical features of potential compounds. The first approaches were frequently based on rule sets [7] with scores awarded to compounds for not failing particular criteria of “drug-likeness”. From the pharmacokinetic perspective, it was proposed to characterize compounds according to absorption, distribution, metabolism, and excretion criteria (ADME) [8]. Further developments have led to refinements of simple rule-based methods into more granular qualitative measures, like Quantitative Estimate for Drug-likeness (QED) [9], which uses a desirability function to compute an optimal score across multiple chemistry-based criteria. Importantly, most of these efforts were not specifically intended only to identify likely toxicity, but to also optimize over a range of relevant properties that can impact efficacy, bioavailability and pharmacokinetics.

A recent evaluation of current methods was performed by [10]. Their work has shown that chemistry-based scoring and rule-based systems have only very modest power to predict clinical trial failures. These methods could not accurately predict clinical trial failure due to drug toxicity if taken in isolation and not combined with additional features. One possible explanation is that these schemes, like Lipinski’s Rule of 5 [11], are now routinely used to screen drugs [12] and compounds at clinical trial stage likely to have already passed such screening. Another part of the explanation could be that these rules do not strongly apply to a very large subset (estimated at 50-80% of all drugs) of “metabolite-like” compounds that can mimic naturally-occurring metabolites and behave in a similar way [13]. And lastly, toxicity-related responses are mostly mediated by drug-protein interactions [14], which may not necessarily have a clear correspondence to molecular structure features.

Given complex nature of drug toxicity prediction problem, chemistry-lead approaches outlined above are just one of the many possible ways to consider it and other studies have explored a wide variety of alternative strategies. Several proposed methods have used various semantic similarity [15] and correlation measures, like known side effect profiles [16, 17] to predict specific side effect labels. An alternative perspective was developed by [18], who used predicted binding to a small number of already known toxicity-related proteins as indicator of risk. Yet another sets of strategies rely on leveraging gene expression [19, 20] and metabolomics profile similarities [21]. Although all of these works have undoubtedly greatly contributed to our understanding of patterns and mechanisms of toxic side effects, not all of these approaches can be used during drug development process because large amounts of *in vivo*, human-specific data can only be safely collected once the risks of the candidate drug are sufficiently understood.

Another complication arises when attempting to integrate or fairly compare approaches, because often scope or prediction goals of different methods are not readily comparable. Frequently, specialized methods are developed for particular classes of compounds [22] or specific, carefully defined scenarios [23, 24]. While most methods measure their success in terms of their ability to predict all possible types of side effects (i.e. both relatively benign to highly dangerous), others [10, 25] consider drug toxicity in terms of drug trail failures or withdrawals from market – a criteria most similar to the one used in this study. Clearly “drug rejection” criteria is not directly comparable with the “side effect prediction” criteria, as in this case the most dangerous side effects are prioritized and “unsafe” category assignment is indirectly affected by the factors like severity and frequency of toxic responses and ability to effectively manage those risks. Although both [10] and [25] studies used comparable criteria, the latter made extensive use of drug annotation (phenotypic, indications and all known adverse reactions). Such data can only be collected once the drug was in use for some and is not available for new compounds, like novel candidate drugs from ClinicalTrials.gov database used in this study.

Drug toxicity is commonly classified into two subtypes: Type A or intrinsic toxicity, which is dose-dependent and related to the primary pharmacological target of the drug, and Type B or idiosyncratic toxicity (IT), which is unpredictable, occurs at frequencies of less than 1 in 5000 cases [26], is not dose dependent and is associated with off-target effects [27]. Although decisions to withdraw a drug from market can be made for a variety of reasons, unacceptable idiosyncratic toxicity is believed to be the main cause [28-31]. Given IT is very rare, it can be missed in smaller test populations used for clinical trials and often is not detectable in animal models [27]. Our analysis indicated that current leading methods developed for the clinical trial success prediction and drug likeness do not perform as well in the case of drugs withdrawn from market (Fig S1). This may indicate that a different perspective or drug properties are needed in order to specifically capture those effects. However, we would like to emphasize that data used to develop these tools and their goals were not optimized for prediction of drug withdrawals from market, and the result reported here is by no means representative of performance of these tools in their intended contexts.

**Figure 1.**
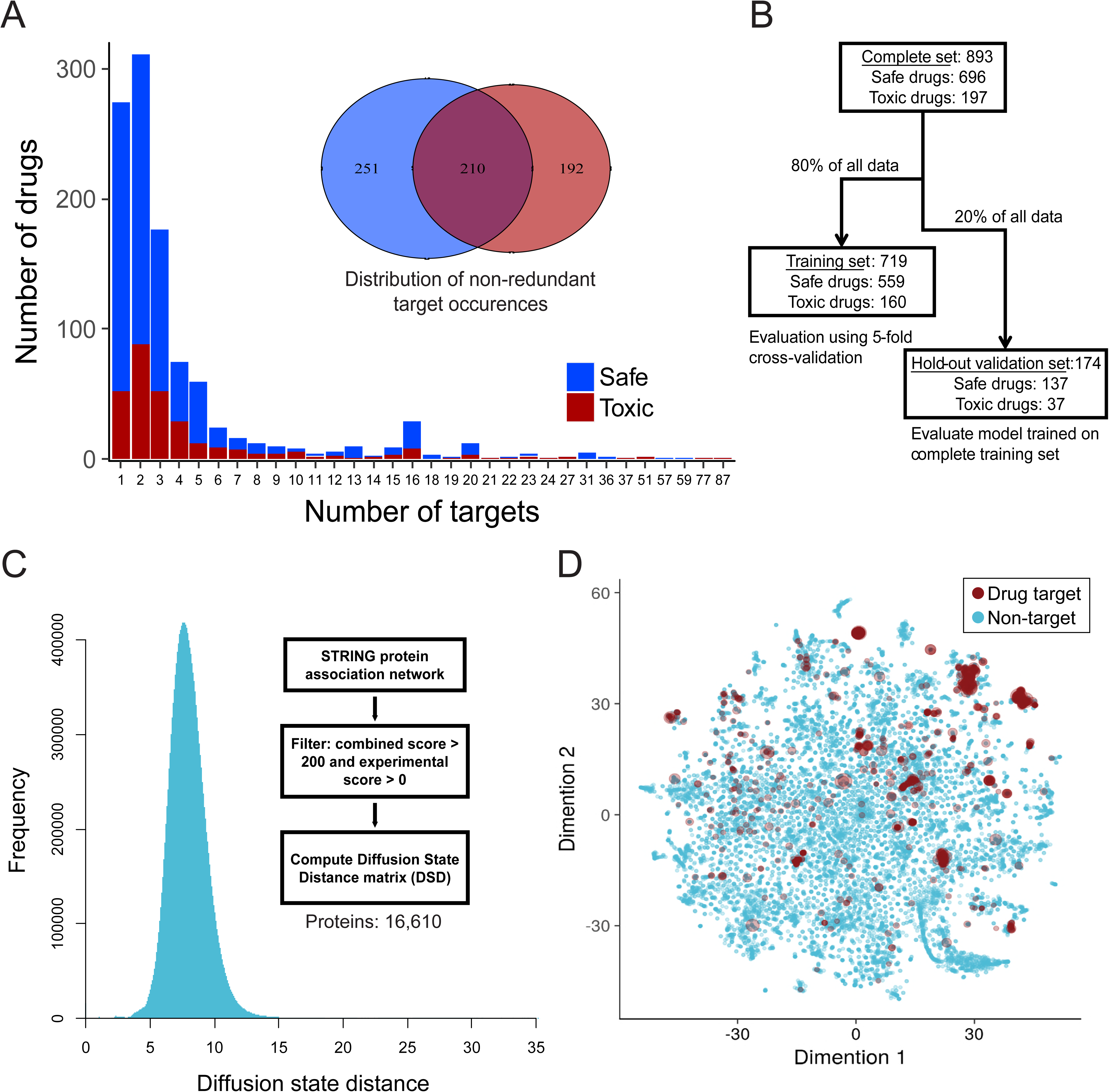
Overview of network, drugs and drug-binding protein data. **(A)** Counts of bound proteins (main target and all off-targets) of each drug in complete datasets. **(B)** Counts of drugs in different subsets used for machine learning model development. **(C)** Overall distribution of diffusion state distances for the main connected component of the network. **(D)** Relative positions of all proteins (n =16,610) in the diffusion state distance space. All pairwise distances were mapped to two dimensions using t-Distributed Stochastic Neighbor Embedding (tSNE) algorithm. Red circles show all drug targets, size of the circle is proportional to the number of different drugs targeting that protein.

Motivated by the importance of drug-protein interactions to drug toxicity mechanisms [14], and the increasing prominence of target-based drug development, in this study we explore the feasibility of developing a computational <u>Target</u>-driven drug T<u>ox</u>icity prediction method (TargeTox). The method uses information about all proteins that can bind a drug (both intended pharmacological targets and off-targets) in combination with machine learning to identify potentially toxic compounds. Importantly, drugs can have both Type A and Type B toxicity risks at the same time and therefore a combination of these factors can lead to a conclusion that particular drug is unsafe. At present, no relevant databases provide structured and comprehensive information about Type A and Type B toxicity risks, however it is generally believed that Type A toxicity is predominantly discovered during clinical trials and Type B – during monitoring and reporting stage, after release to market [27]. For these reasons when designing a training dataset we aimed to include examples for both cases, though in our downstream analysis we place particular emphasis on confirming performance for idiosyncratic toxicity cases. Although we aim to predict likely risk toxicity of both types, current implementation is not designed to directly identify the type.

One particular challenge in incorporation of drug-binding protein data is the sparse nature of the dataset, where each drug will only bind a relatively small set of all possible proteins and this number will differ between drugs. At the same time, given that the set of confirmed toxic drugs satisfying our criteria is small, large numbers of bound proteins and most of interactions will occur only once in the entire dataset. To address this, we propose to leverage guilt-by-association principle in combination with biological network context of these proteins. Since it was previously reported that target proximity in the network corresponds to the similarity of drug side effects [32], we hypothesized that severe toxicity-related responses could be localized to particular regions of the biological network. Here, network is represented by a distance matrix of all constituent proteins. As part of this study we have found that both simpler and more sophisticated network distance measures can be used with this approach, though based on our evaluations diffusion state distance (DSD) [33] was chosen as the marginally best-performing metric. The position of each drug-specific set of bound proteins is approximately encoded by distances to a small number of reference proteins. This interpretation of the data allows all observations to be meaningfully used, including cases where single instances of drug-binding proteins are found in training or evaluation datasets. Although concepts of biological network diffusion have been explored in other contexts, e.g. [34], the distinguishing and novel feature of our approach is in direct use of a machine learning classifier to “map out” areas of the network during the training process, which means effects of other covariates can also be taken into account in conjunction with network-based location data. At the same time, this method can also be used to reduce dimensionality and convert data from sparse to dense representation.

## Results

### Drug-binding proteins tend to be non-uniformally distributed in the network

Information for all drug-binding proteins in our reference set was acquired from DrugBank and ChEMBL databases. Although there was a substantial number of drugs with a single target (Fig 1A), the majority of drugs interacted with more than one protein. The number of bound proteins was also smaller than the number of drugs, and about 47% of all these proteins were found in both toxic and safe subsets. To explore the overall distribution of drug-binding proteins in the context of human interactome they were combined with a protein association network from STRING database, which was transformed into a diffusion state distance (DSD) matrix to do this analysis. Overall DSD distribution for all proteins in the main connected component of the network was largely consistent with what was previously reported by [33] for yeast protein-protein interaction network (Fig 1C). Generated distribution had a relatively smooth central part with a long right tail. To visually explore possible location patterns of bound proteins, we mapped the complete DSD distance matrix into two dimensions (Fig 1D) using t-Distributed Stochastic Neighbor Embedding (tSNE) algorithm [35]. Although a minority of bound proteins appeared to be dispersed throughout the network, most tended to be co-located in a couple of distinctive groups. On average, bound proteins of the same drug tended to be significantly closer together than random samples of the same size, though there was no difference between average distances of toxic and safe drug-interacting protein sets (Fig 2A) and the same pattern was observed when only a subset of drugs withdrawn from market was considered (Fig 2B). The overall proximity of the proteins binding the same drug suggested a possibility that these sets may be represented more compactly by network locations in order to reduce sparseness and dimensionality of the data while minimizing the loss of useful information.

**Figure 2.**
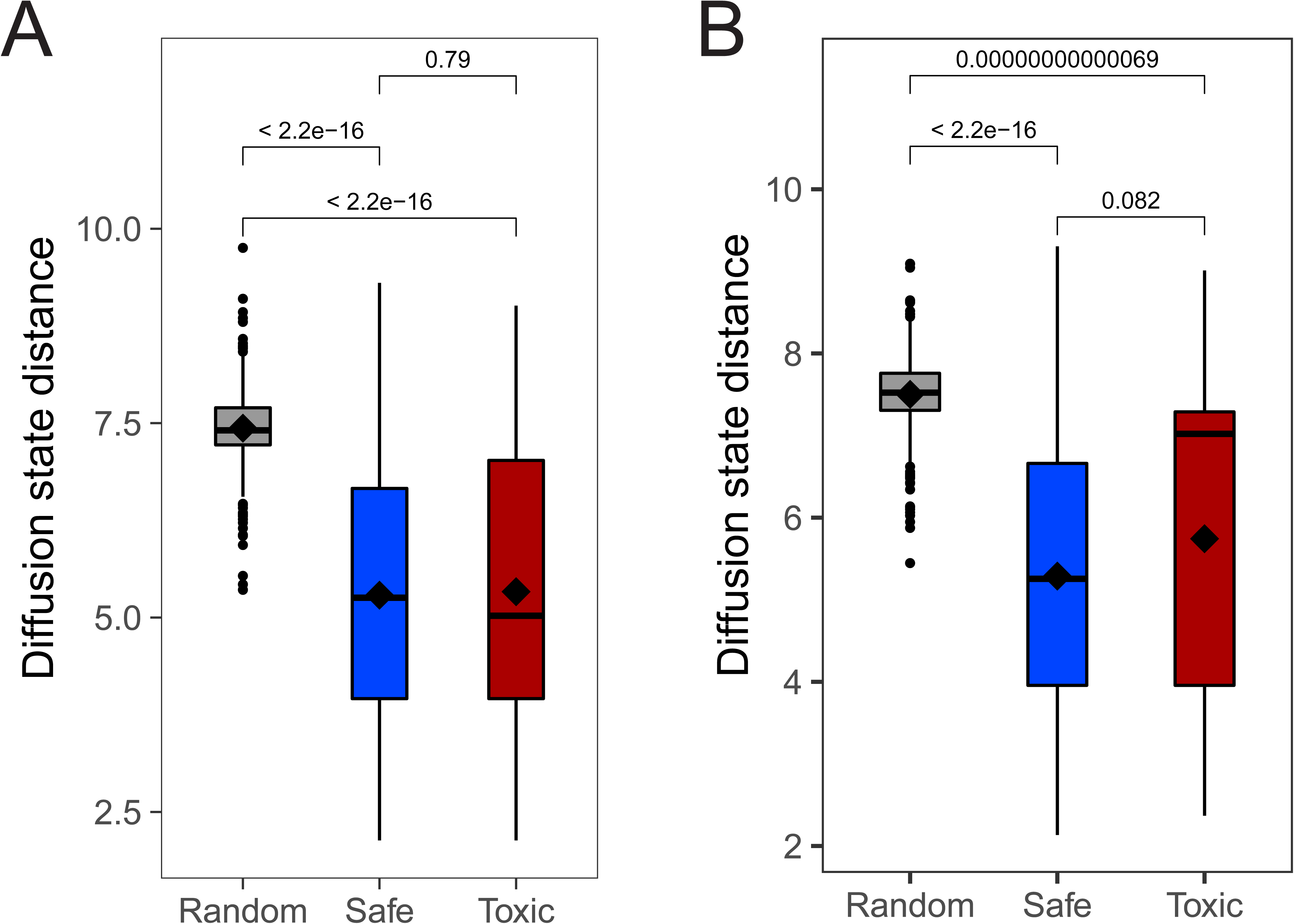
Comparison of distances between proteins binding the same drug and a random sample. **(A)** Diffusion state distances between proteins bound by the same drug for all drugs included in the study. **(B)** The same comparison for the drugs withdrawn from market for reasons of toxicity. In both cases the significance was computed using Wilcoxon signed ranks test.

### Computational model for prediction of dangerous drug toxicity

To facilitate accurate identification of potentially toxic drug candidates, we have developed a Biological Network <u>Target</u>-based Drug <u>Tox</u>icity Risk Prediction method (TargeTox). The method aims to leverage the guilt-by-association principle, according to which entities close to each other in biological networks tend to share functional roles. Distance between nodes in biological networks can be quantified using a variety of different methods and we have evaluated several strategies ranging from very simple, like shortest path, to novel and advanced approaches, like Mashup [36] method that integrates diffusion-based distances across multiple ‘omics networks. By interpreting a network as a set of pairwise distances, biological functions and phenotypes can be associated with areas of the network rather than just individual nodes, and their location can be efficiently summarized with respect to a few reference points. Once the network location data has been put into this form and combined with relevant covariates, a machine learning classifier was trained on the combined dataset.

In principle, this strategy can be used in combination with any modern classifier that has some form of regularization capabilities and can handle non-linear relationships, e.g. certain SVM variants or deep neural networks. However, in this case gradient boosted classifier tree ensemble model (GBM) was chosen for the following two reasons. First was the small numbers of positive (toxic) drugs in our training dataset, which meant that comparatively less hyper-parameter tuning required by GBM was considered to be very helpful for mitigation of the overfitting risk. Second reason was the presence of missing values in our data, which GBM can handle without the need for prior imputation, thereby greatly simplifying both development and any possible future applications of our method.

To control for the risk of over-fitting our model, the available data was split into a training set (80% of all drugs, Fig 1B) and a hold-out validation set (20%). The performance of different design strategies and hyper-parameter configurations was evaluated on the training set using five-fold cross-validation, then a model was trained on the complete training set and evaluated on the remaining twenty percent of the data. We have evaluated the following strategies for measuring network distances: shortest path, discretized shortest path (1 if less than length 3, 0 otherwise), diffusion state distance [33] and Mashup [36] methods. Our evaluation results showed that generally all of the tested measures can to some extent be used in combination with our method, however, DSD-based implementation had the best performance by about 2-3% of ROC AUC (Fig S2 A,B,E,F). Additionally, we have evaluated two different ways of summarizing drug-binding protein information: (1) using a medoid protein for a set of all proteins binding a particular drug and (2) using distance to all other proteins in the network rather than choosing a few reference proteins. The first strategy had achieved 69.69% ROC AUC (Fig S2 D). The second strategy was the second-best performing of all tested configurations (ROC AUC of 72.79%, Fig S2 C), however at a great cost of time needed to train the model. For the latter strategy, we also observed that in actuality only some of all points were used in the trained model, as GBM performs feature selection during training. The best strategy (ROC AUC of 73.35%) was to use a small number of reference points with a DSD metric, and for each of them take a distance to the closest protein bound by a given drug. On the training subset, at optimal trade-off point, this strategy achieved sensitivity and specificity of 74.7 and 65.8, respectively. On hold-out test set ROC AUC for the chosen optimal version was found to be 71.30%. Feature importance analysis done on the final version of the model (Fig 3 D) indicated that in aggregate network-based features where the most important category accounting for half of all importance, while functional impact feature was the most important single feature. No features were discarded as a result of feature selection done by the algorithm during training.

**Figure 3.**
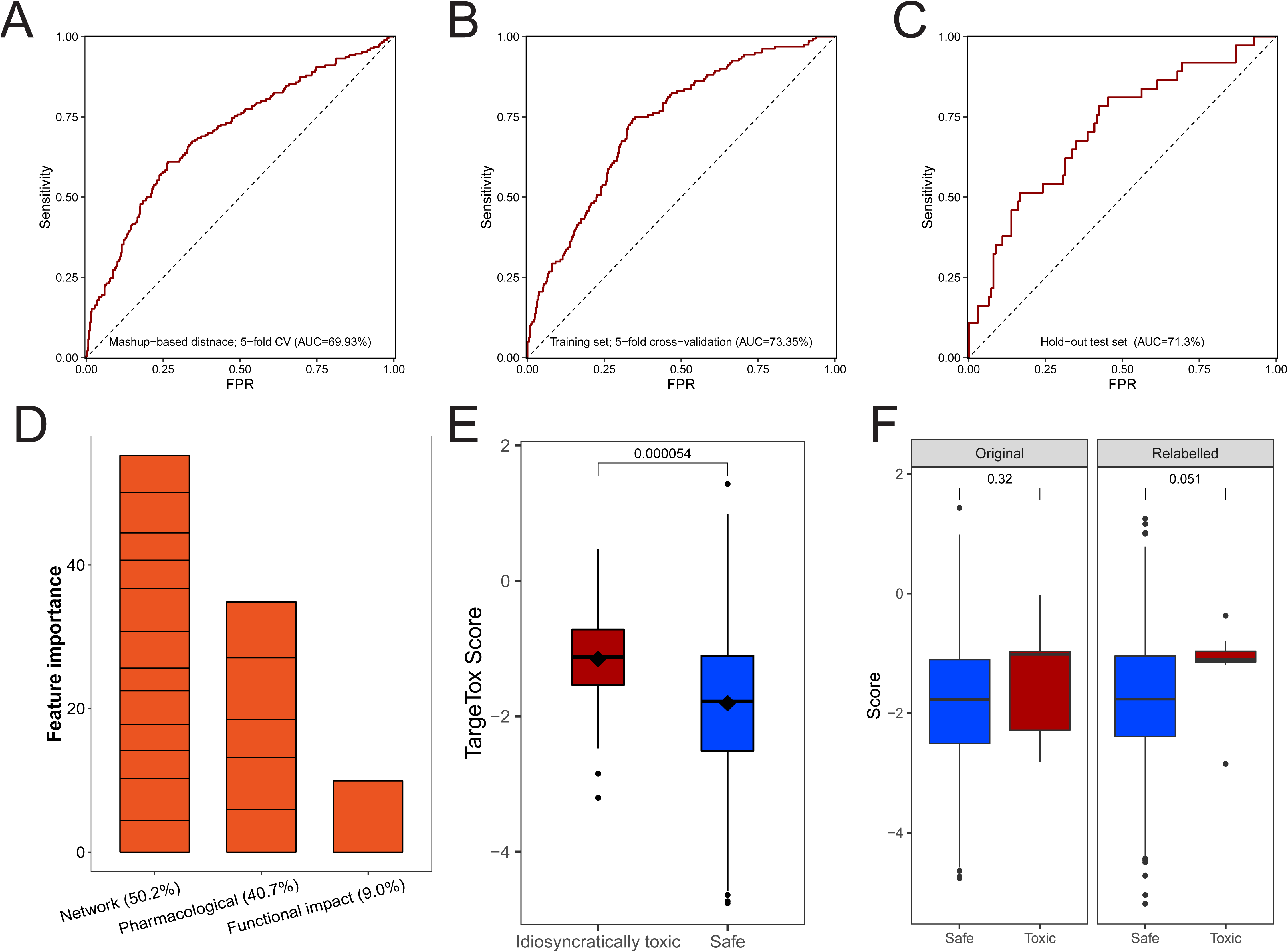
Performance benchmarks and model interpretation. Receiver-operator characteristic (ROC) curves for different model variants. **(A)** Mashup-based distance version evaluated on the training set using 5-fold cross-validation (CV) (n=719); **(B)** Diffusion state distance version evaluated on the training set using 5-fold CV (n=719); **(C)** Diffusion state distance version validated on the hold-out set (n=174). **(D)** Contribution of different features to the model measured in feature importance. **(E)** Comparison of scores returned by the model for the idiosyncratically toxic (n=38) and safe subsets (n=696) computed using final model and leave-one-out cross-validation. **(F)** Comparison of scores for toxic drugs linked to HLA-mediated toxicity (n=9) computed using leave-one-out cross-validation. Original sub-plot shows scores if curation-based toxicity annotation was used, relabelled sub-plot shows scores when all relevant drugs are relabeled as toxic. Significance was computed using Wilcoxon signed ranks test.

### Evaluation of ability to predict idiosyncratic toxicity

To investigate potential of the method to identify idiosyncratically-toxic drugs, we have created two relevant subsets. The first had 38 drugs from our dataset that have been specifically identified as idiosyncratically toxic in the literature. The second subset had 9 drugs specifically associated with the HLA-mediated toxicity [37]. HLA-mediated toxicity is one of the prominent and relatively well-studied examples of idiosyncratic toxicity. Therefore these drugs could be used to explore the ability of our approach to identify potential common toxicity mechanism for a group of drugs.

To explore the performance for the more general set of 38 idiosyncratic toxic drugs we have done a leave-one-out cross-validation and compared the scores of the 38 drugs to those in the safe subset. The scores were consistently and significantly higher (i.e. predicted to be more toxic) for this subset (Fig 3 E). Then, we have done the same comparison for the scores from PrOCTOR and weighted QED methods, but no significant differences were detected for either method (Fig S3). To explore whether our chosen features could capture patterns specific to idiosyncratic subset, a more detailed feature attribution analysis was done using SHAP (Shapley additive explanation) values methodology for gradient boosted tree ensembles [38]. After computing feature-specific SHAP values for each drug, we have compared idiosyncratically toxic subset with drugs where toxicity was identified during clinical trials. For the latter subset, we have verified that idiosyncratic toxicity was not reported as the main cause of clinical trial termination in the corresponding entry of the ClinicalTrails.gov database. Additionally, to the best of our ability, we have checked for other factors that could bias these results, including over-representation of particular drug classes or indications. Each pair of SHAP value distributions was compared using Wilcoxon signed ranks test that identified 7 significant differences at 5%, of which one was also significant at 1% level (Table 1). These results suggest that substantial number of features identified as particularly important for Type B versus Type A toxicity are distinctive and these differences were captured by our method design.

In a second, more specific example we have identified 9 drugs known to be idiosyncratically toxic via an HLA-mediated mechanism. Of all the drugs in this category, only 3 were already categorized as toxic according to our chosen criteria (clinical trial or market withdrawal for reasons of toxicity). One possible explanation could be that given that this particular mechanism of idiosyncratic toxicity is very well researched, effective strategies exist (e.g. known risk alleles, populations and treatment regimens) to manage this risk and allow most of these drugs to be used relatively safely. Similarly, leave-one-out cross validation (Fig 3 F) using original safe/toxic assignment did not indicate that these drugs were significantly more toxic that the main “safe” category. Next, in order to further explore the potential of our method to detect common toxicity mechanism of this group of drugs, we have done an additional leave-one-out validation where all of them were relabeled as “toxic”. This change did cause an increase in the predicted toxicity score for most members of the set, however overall this difference was not found to be significant at a 5% level (Fig 3 F).

### Independent validation using side effect annotation

A secondary validation was done using side effect annotation from the OFFSIDES database [39], from which we selected drugs not present in either training or hold-out subsets. After pre-processing, the validation dataset contained 339 drugs. Given the wide scope and diversity of possible side effects, many of which are not considered severe enough to preclude the use of a drug, we have selected fourteen toxicity-related categories commonly associated with failed drugs, including “cardiotoxicity”, “hepatotoxicity” and “toxic shock”. Predicted scores of drugs in these subsets were compared with a set of 120 compounds that did not have any of these annotations (Table 2). The average score of these major toxicity-associated categories was higher than average of the unannotated set, and the difference was significant in all individual cases except for “nephropathy toxic” and “mitochondrial toxicity” categories. Likewise, the overall difference of pooled set had a significantly higher average score (Fig 4). These results reaffirm the particular relevance of the proposed method for identification of high-risk drugs of these types.

**Figure 4.**
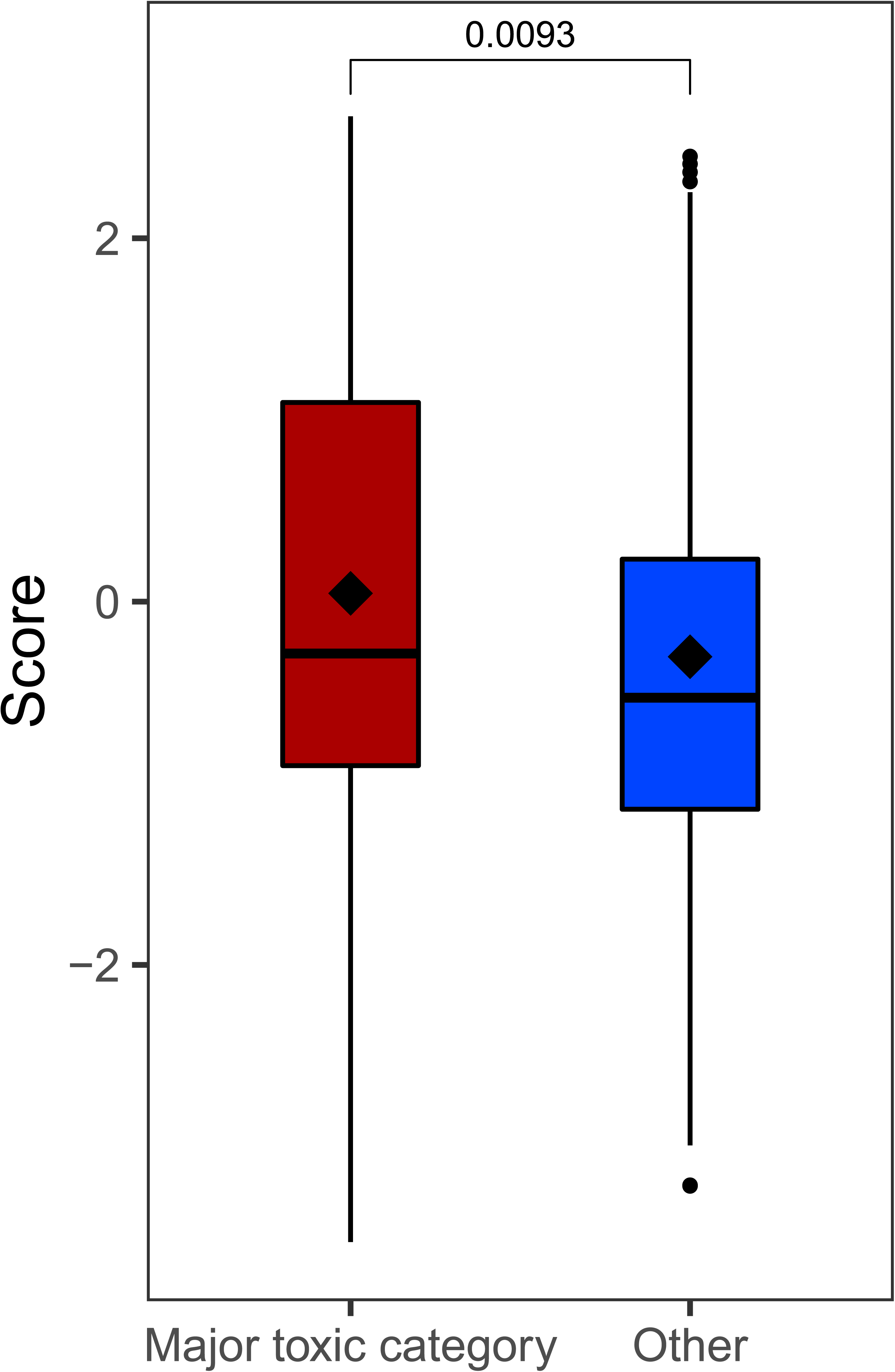
Comparison of scores for annotation-base toxicity categories. Major toxic category contains all drugs with toxicity-related annotations from OFFSIDES database (n=257), safe category contains all drugs without such annotations (n=120). Significance was computed using Wilcoxon signed ranks test.

### Model interpretation

Although gradient-boosted tree ensemble methods are very powerful and flexible, the complexity of generated models makes them challenging to interpret directly. An additional complication arising from the chosen design of network-based features is that by itself individual importance of a reference protein feature may not directly identify bound proteins associated with toxicity risk, rather location of relevant proteins can be defined by a higher-order interaction of several such features. Nevertheless, this information is captured by the model and can be extracted in order to profile potential toxicity risk of different proteins.

In order to extract this overall “toxicity risk map” from the model, we have created a simulated dataset of single-target drugs for each of the proteins in the “druggable genome” list from the work of [40]. Most of this set (4019 proteins) could be mapped to the main connected component of the STRING protein association network. Notably, the highest score achieved by a simulated single-target drug was 45% lower than the top score in a real dataset, indicating that highest predicted toxicity risks are due combined effect of multiple causal proteins. Despite this important difference, these results could still be useful for interpreting the behavior of the technically “black-box” model and extraction of informative insights. Distributions of the scores assigned to these proteins by TargeTox were visualised to check for the presence of the coherent structure. Again, this was done with the aid of t-SNE to project the positions relative to the twelve reference points into two dimensions. These results (Fig 5) showed that bound proteins predicted to have a higher risk are concentrated in several hot spots. Top 10% of these predictions were separately clustered to identify possible high toxicity risk sub-groups. Clustering suggested the presence of 8 subgroups, four of which (1-3 and 6) corresponded to quite compact and distinctive neighborhoods suggested by the t-SNE algorithm and four others were distributed over wider areas.

**Figure 5.**
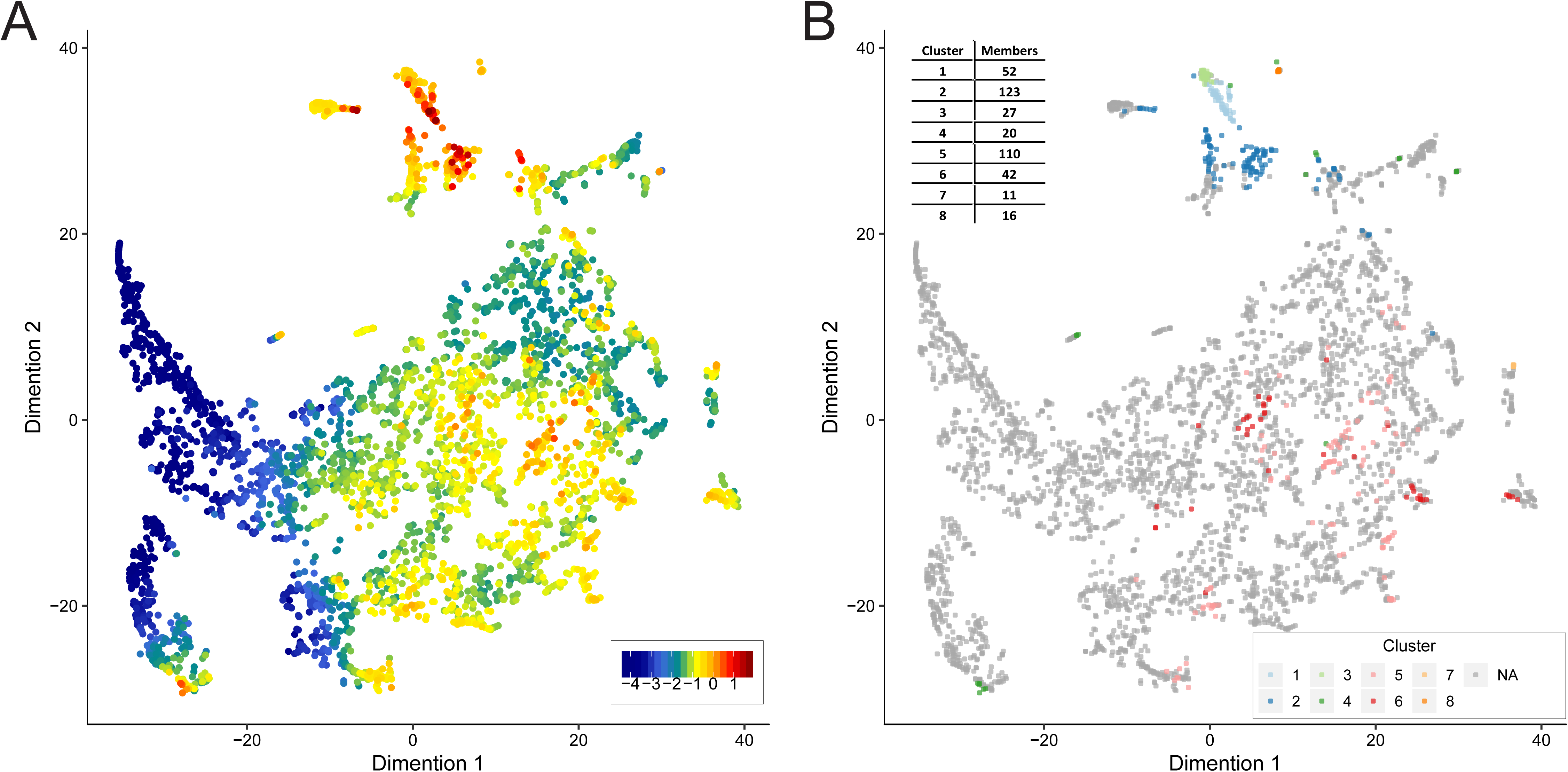
Druggable proteome annotation with TargeTox method. Two panels show druggable proteome (n=4019) in diffusion state distance space computed from STRING protein association network and mapped to two dimensions using t-SNE method. **(A)** protein nodes colored according to TaregeTox score (red = highest risk, blue = lowest risk). **(B)** location of distinctive sub-groups with highest risk identified using clustering of top 10% of all proteins by TargeTox score.

Common functional roles of these protein groups were identified using functional enrichment analysis (Fisher’s exact test with False Discovery Rate correction) with respect to Biological Process aspect of the Gene Ontology. Overall, most common recurring processes included signaling and protein phosphorylation, with multiple significant hits across all clusters, with the highest fraction in Cluster 1 (94.12% of all proteins, p = 0.002). The highest predicted toxicity risk scores were particularly concentrated in Clusters 1 through 3, which were also placed close together by the t-SNE algorithm, suggesting similar protein-protein interaction context. Some notable potentially relevant functions included immune-related processes (clusters 1, 2, 5 and 6, multiple). Disruption of immune system processes frequently underlies toxic side-effects [41]. Clusters 1 and 3 were enriched for peptidyl-tyrosine phosphorylation (86.3% of all members, p = 4.02×10^-30^ in Cluster 1 and 70% in Cluster 3, p = 0.002) and, in Cluster 1, also positive regulation of JAK-STAT cascade (13.73%, p =2.74×10^-4^). Tyrosine kinases are prominently linked to idiosyncratic toxic side-effects, including cardiotoxicity [42], whereas *JAK-STAT* pathway is important for different aspects of neurologic toxicity [43]. Cluster 5 was significantly enriched for response to toxin (15.38%, p=0.03). Cluster 4 had high number of G-protein coupled receptor signaling pathway members (60%, p = 0.01). Apoptotic process, believed to play an important role in drug-induced hepatotoxicity [44] and cardiotoxicity [45], was the largest enriched category in Cluster 7 (63.62%, p=0.01). No significant GO term enrichment for any functions was identified in Cluster 8. Full results of this analysis in the form gene annotations, their cluster assignments and GO biological process enrichment are provided in the supplementary (Tables S1 and S2).

In terms of individual proteins, the highest toxicity-scoring predictions were concentrated in Clusters 1-3. Top predictions featured several proteins identified as promising anti-cancer drug targets. The highest-ranked protein with the score of 1.77 was FGFR2, a tyrosine kinase and an important oncogene. In particular, this protein binds AZD4547 candidate drug, clinical trials of which have reported a number of serious toxicity incidents [46]. The third highest-scoring protein TLR4 is suggested to play an important role chemotherapy-induced gut toxicity [47] and nephrotoxicity [48]. Among other proteins in the top ten were AKT1, KIT, JAK2 and LYN. AKT1 is an potassium channel protein, inhibition of which was found to be linked to liver injury and development of hepatocellular carcinoma in animal models, with implications for human clinical trials currently in progress being very likely [49]. Proteins KIT, JAK2 and LYN are all members of the tyrosine kinase family that both has many promising drug targets while also being associated with serious toxicities, both on-target [50] and unexpected [51] and, more specifically, idiosyncratic hepatotoxicity [52]. One notable example somewhat further down the list was PTGS2 (COX2), which was rated in the top 2% for likely toxicity and is known to be the key protein target leading to high-profile withdrawal of Vioxx drug due to doubling of heart attack risk [53].

## Discussion

One principle obstacle to the development of computational predictive approaches is the sheer complexity of the interplay between factors that determine whether a drug is considered to have an unacceptable level of toxicity. (Fig 6). The fundamental trade-off at the core of this decision is striking the right balance between efficacy and safety [54], both of which are evaluated with respect to the severity of the disease to be treated. Each of these factors may be subject to considerable variation. Only a small number of people might experience a toxic side effect and it may have different degrees of severity and, in the case of idiosyncratic responses, exact underlying causes can be particularly complex [55]. And lastly, extrinsic factors like cost, availability of alternative treatments, and ability to predict or manage risks also inform the final decision [56]. Although ideally it is very desirable to directly incorporate the effects of these factors into a predictive toxicity model, at present such data is still not systematically collected at the necessary level of detail. Not being able to accurately model these effects is an important factor limiting the accuracy of computational drug screening approaches, but structured data collection efforts by initiatives like <u>ClinicalTrials.gov</u> are likely to address this data availability problem in the near-future.

**Figure 6.**
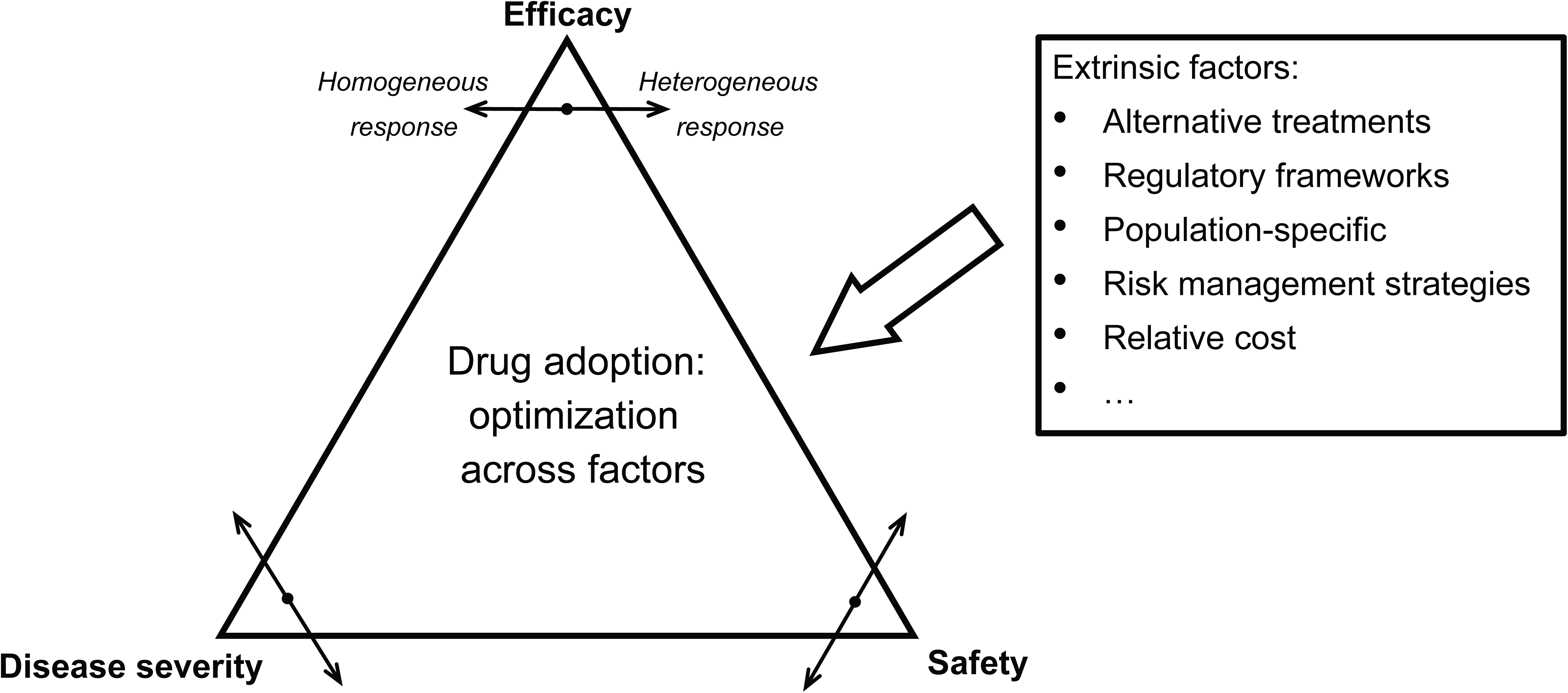
Factors affecting drug safety decision-making.

Another limitation is the actual number of failed drug observations that are currently available in the public domain. Very large number of features may need to be included in the model to adequately capture underlying complexity, which in turn would necessitate a large number of observations (example drugs) to accurately profile their effects. One way to deal with this issue could be to mirror the drug development stages in separate steps of computational screening pipeline. In early stages, the drug development process primarily focuses on chemical features of screened compounds and their pharmacokinetic features, then biomedical and clinical contexts are considered during laboratory and clinical trial testing. Computational models can be specialized to achieve optimal performance for each of these stages and combined to form a sequential filter, e.g. approaches like QED and ADME can be used as a first step, then tools like PrOCTOR to identify compounds likely to fail during clinical trials and, lastly methods like ours to identify the remaining problematic compounds. The only essential input required for TargeTox is identity of proteins bound by a particular drug, while optional inputs, which may be missing, are the three Boolean values for possible routes of administration and two numeric values for lower and upper protein binding. All of the other features, which are actually used by the trained model, are computed from the supplied list of bound proteins and the implementation released with this paper can perform this part of the analysis automatically.

Our method is particularly dependent on knowledge of proteins binding specific drug, as this data is necessary to compute both the network-based and functional impact features. Idiosyncratic toxicity is often mediated by the effect on off-target proteins of particular drugs [57]. Information about all possible bound proteins can often be incomplete, which can limit the effectiveness of the proposed method. Some resources offer computationally-derived predictions of bound proteins [58], though use of such information would necessitate striking a right balance between true and false positives. Another importance aspect not currently considered by our model is the metabolism of the drug, which can generate toxic secondary compounds that may result in idiosyncratic toxicity and can also have their own sets of protein interactions [59]. Development of effective strategies for incorporating this wider body of knowledge can lead to further improvements of TargeTox and other similar methods.

Another important means of further progress could be in better utilization of other types of biological network data. This particular consideration was partially explored by considering distances derived from an integrated set of networks using Mashup method. Although in this particular case we have found that a single network of experimentally confirmed protein-protein interactions was marginally better, it is very likely that the results better results could be obtained using different combinations of networks or different link reliability thresholds. Although we have not been able to comprehensively explore all of these options as part of this initial study, we believe that more in-depth evaluation of integrative methods like Mashup merits further investigation. Additionally, incorporation of data from cell culture profiling studies offered by the Connectivity Map (CMAP) [60] and the more recent Library of Integrated Network-based Cellular Signatures (LINCS)[61] could be another way of more fully capturing complexity of drug responses. The potential of combining such data with network-based approaches was recently demonstrated by the SynGeNet method [62], which successfully predicted genotype-specific drugs for melanoma.

In this work, we were particularly interested in exploring the problem of idiosyncratic toxicity, i.e. where the toxic effect is only manifested rarely and therefore may be missed during clinical trials. We were able to confirm that existing methods were not as effective in identifying these drugs. We have presented an example showing that our method can improve on the performance of existing methods specifically for those drugs. By applying TargeTox in a speculative way, we were also able to generate toxicity risk annotation for druggable proteome. This follow-up analysis suggested that bound proteins associated with predicted toxicity risk are concentrated in highly specific areas of human interactome and tend to have immune system and signaling-related functions. An additional insight was that highest toxicity risk scores were only predicted by our model when a drug had several targets. This suggests that burden of multiple drug-protein interactions on particularly susceptible regions of the networks could be a plausible hypothesis for explaining most severe cases of drug toxicity.

To facilitate applications of TargeTox we have made the trained model, supporting data, and necessary code available in a dedicated GitHub repository (https://github.com/artem-lysenko/TargeTox). Given that the only essential input for TargeTox is the identity of bound proteins for each drug, the method has particularly good synergy with the currently dominant target-driven paradigm of drug development. We believe the method will be particularly useful for identification of idiosyncratically toxic drugs during a computational screening of drug compounds as well as for the prior knowledge-directed design of combinations that minimize toxicity risk.

## Methods

### Dataset construction

Reference dataset was based on three resources: DrugBank [63] for drugs currently in use, <u>ClinicalTrials.gov</u> [64] for drugs that failed clinical trials and supplementary data from [4], which compiled a comprehensive list of drugs that were withdrawn from market between 1950 and 2016. The latter two resources were filtered manually to only keep the drugs that have failed for toxicity-related reasons. DrugBank dataset was also filtered to remove all antineoplastic drugs, as those are expected to have relatively high toxicity to be considered sufficiently similar to the “safe” drugs for other diseases. To resolve any naming ambiguities, all drug names were mapped to ChEMBL identifiers using DrugBank database and manual curation. Duplicates were removed to retain only one entry, with precedence given to the “toxic” class subset. Then, the ChEMBL database [65] was used to obtain the chemical structure information (SMILES strings) and bound proteins (both main pharmacological target(s) as well as any off-targets) for each drug. All entries where no complete information was available were discarded at this stage. This resulted in a set of 696 compounds in the “safe” category and 197 compounds in the “toxic” category. The ChEMBL and DrugBank databases were also used to put together pharmacological covariate data, specifically route of administration (oral, parenteral and topical) as well as lower and upper estimates for blood plasma protein binding, though missing values were allowed for these features.

Data for proteins bound by each drug was integrated with a protein association network from STRING database [66]. In order to control for false-positive edges, we only used experimentally-confirmed interactions with a combined score of at least 200. To ensure consistent distribution of distances, only main connected component of this network containing 16,610 proteins was used for all of the analysis. Information about the biological function of the bound proteins was acquired from Gene Ontology [67] annotation database and annotation of drugs with side-effects – from OFFSIDES database [39]. Additional literature curation was done to identify a subset of drugs with reported idiosyncratic toxicity, which is provided in supplementary (Table S3). HLA-mediated toxicity drugs were identified based on the list from [37].

### Computation of candidate predictive features

Chemical structure was used to compute drug properties using the ChemmineR package [68] applied as specified in PrOCTOR analysis script [10]. Additionally, we ran all drugs in our dataset through PrOCTOR to obtain the PrOCTOR score and weighted quantitative estimate of drug-likeness (wQED) [9].

Several different network distance metric were evaluated for inclusion in our model. Simpler metric considered were the shortest path length in STRING protein-protein interaction graph and the discretized shortest path where a value 1 was assigned if length was less than 3 and 0 otherwise. Of the more advanced measures we have considered Mashup, which computes distance over an integrated set of multiple biological networks, and Diffusion State Distance algorithm, a distance measure based on random walks. In the case of Mashup, we have used the pre-computed matrix of vectors for STRING networks made available by the authors of the algorithm. The matrix was transformed to distance matrix by computing cosine distances between all pairs of vectors. Cosine distance was chosen because it was suggested as the most appropriate one in the original Mashup method paper.

In the latter case, the network was transformed into a symmetrical Diffusion State Distance (DSD) as described in the work of [33], using the following formula:
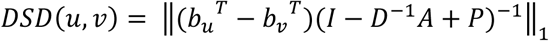

where *D* is the diagonal degree matrix, *P* is the constant matrix of the steady-state distribution and *b_u_*, *b_v_* are the basis vectors for respective nodes. The DSD metric allows the fine-grained mapping of all drug-binding proteins into a network-predicated topological space. Using this distance metric we were able to compare the relative distributions of different bound proteins sets. As we have found that proteins that bind to the same drug tended to be located close to each other in the network, the position of the set can be approximated by the position of its convex hull with respect to a few reference nodes. This transformation summarizes the biological network location of any set of possible bound proteins in the same small number of variables -- regardless of its original size.

Next, we have explored several possible designs for the network-based features. Options considered were representing each drug by a network-based medoid for all bound proteins of a particular drug and using a full set of distances between closest bound protein and each other node in the network. Given that the latter, most promising strategy had considerable computational costs, we have explored how the number of reference points can be reduced. Specifically, the reference nodes were chosen to be most representative with respect to the set of all drug-binding proteins, with the rationale being that this proximity will serve to reduce possible noise and errors due to unavoidable missing or spurious edges in the protein association network. Candidate reference nodes were identified by computing enrichment for drug binding proteins in a fixed-distance neighborhood around each node in the network (Fig 7). To reduce redundancy, all significantly enriched neighborhoods were clustered using hierarchical clustering to group them into the desired number of distinctive groups. Finally, the representative central node of the densest neighborhood in each cluster was chosen as a reference node. Three free parameters of this approach (neighborhood-defining distance, enrichment cut-off and number of clusters) were optimized using grid search.

**Figure 7.**
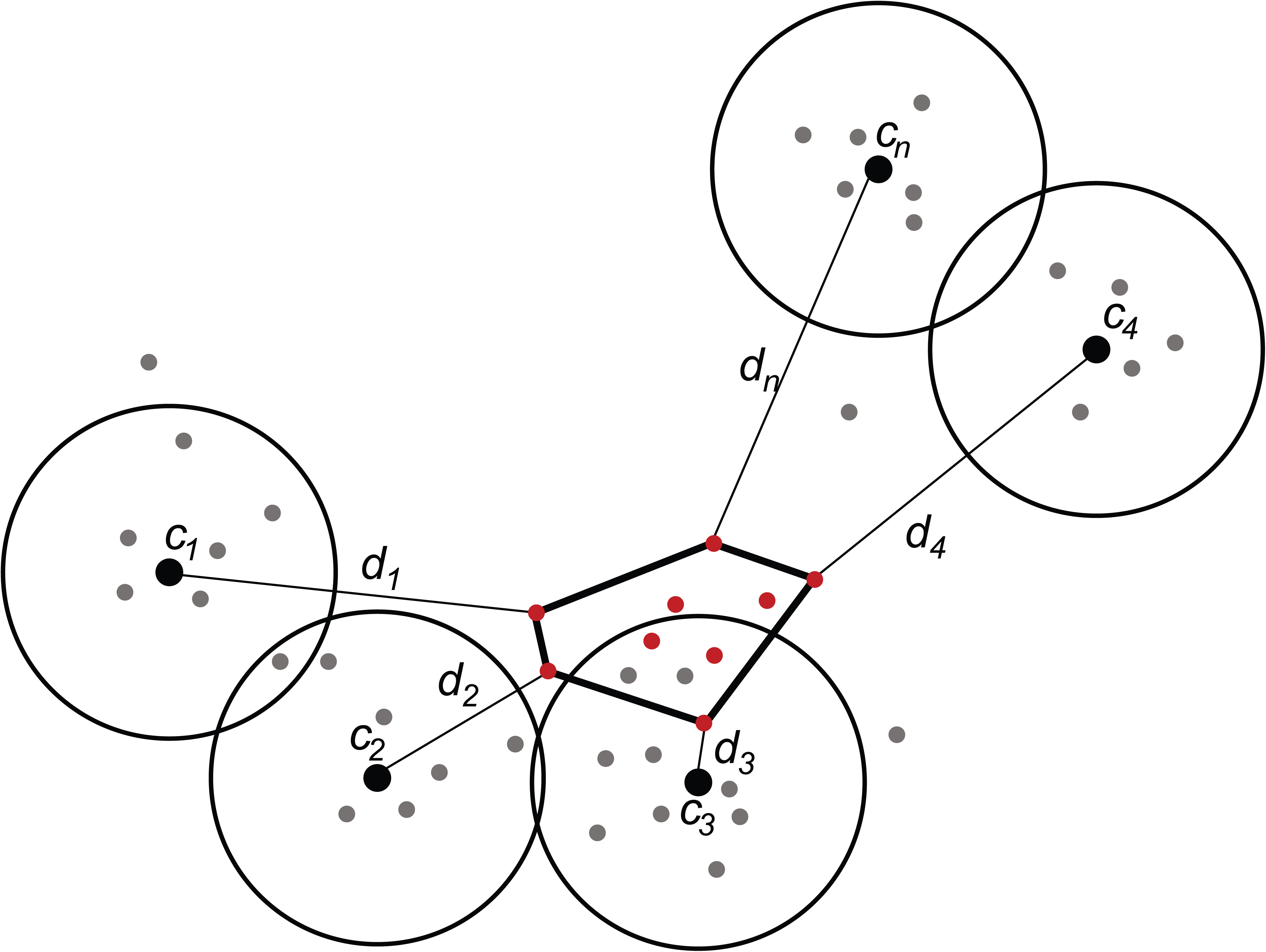
Conceptual schematic of the network-based feature design. All protein nodes are mapped into a diffusion state distance space (filled dots). A set of reference nodes (c1-cn; black dots) are chosen in the areas with dense concentration of known drug targets by computing enrichment in a fixed-distance area around of each candidate node. The redundancy is then removed using hierarchical clustering of all qualifying candidates. The network distance-based features are defined as a distance (d) between a reference node and the closest target for a given drug (red dots).

Functional impact score (FI) metric was derived from the Gene Ontology biological process (BP) annotations. For the purposes of this analysis annotations to each term also inherit annotation to all of its ancestor terms. Using a complete set of all human annotations, information content was computed for each of the individual terms:
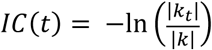

where *t* is a given GO term and *k* is an instance of annotation (unique entity-term pair). Then, functional impact score is defined as follows:
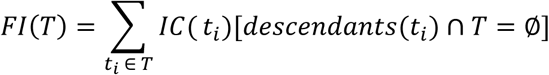

where *T* is a set of all annotation terms for a set of particular drug-binding proteins. The rationale behind the design of this feature is that a drug is expected to have an impact on some biological processes as part of its intended mechanism of action. If this impact is focused, there will be few other processes affected, so the score will be relatively low. On the other hand, if a drug also affects some off-target processes or interacts with a critical protein contributing to multiple processes FI score will be high. Information content serves to achieve even further granularity of the measure, as it is low for generic functions that are relatively common and high for specialized functions where there is little redundancy.

### Classifier training and evaluation

Here we describe basic notations for training and evaluating our proposed model. Let *χ* be a set of *n* samples in a *d*-dimensional feature space which is split into training set *χ_tr_* and test set *χ_ts_*; i.e., *χ* = *χ_tr_* ∪ *χ_ts_*. Let *Ω* = { *ω_i_*: *i* = 1,2, … *c*} be the finite set of *c* class labels and *ω_i_* is the class label of *i*th class. To preserve the classes, the training and test sets are subdivided into *c* disjoint subsets *χ_tr_* = *χ_tr_*_1_ ∪ *χ_tr_*_2_ … ∪ *χ_trc_* and *χ_ts_* = *χ_ts_*_1_ ∪ *χ_ts_*_2_ … ∪ *χ_tsc_*, respectively, where *χ_trj_* ⊂ *χ_tr_* and *χ_tsj_* ⊂ *χ_ts_*. Further, it can be noted that each subset *χ_trj_* or *χ_tsj_* has class a label *ω_j_*. Let *n_trj_* and *n_tsj_* be the number of samples in *χ_trj_* and *χ_tsj_* (of class *ω_j_*) such that
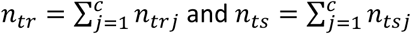

The feature vectors of *χ_tr_* and *χ_ts_* can be depicted as
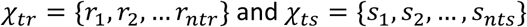

To perform a robust evaluation of the model, we split up our data into training subset *χ_tr_* and validation subset *χ_ts_* at *n_tr_* : *n_ts_* = 80:20 ratio while preserving the ratio of the two classes in our dataset; i.e., *n_tr_*_1_/*n_tr_*_2_ ≈ *n_ts_*_1_/*n_ts_*_2_. During development, evaluation was done using five-fold cross-validation approach using only the compounds in the training set and the final evaluation was done using the holdout set.

To train our model we have chosen to use a gradient boosting algorithm. The objective of gradient boosting algorithms is to find an approximation 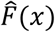 of a function *F*(*x*) such that the expected value of a loss function *L*(˙) is minimum [69]; i.e.,
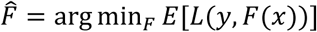

where *y* is the class-label of a feature vector *x*, and *E*[˙] is the expectation function. Gradient-boosting is generally used with decision trees *h*(*x*). The *t*-th step gradient-boosting with decision trees *h_t_*(*x*) is updated in Friedman’s algorithm as
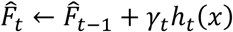

where step-size *γ_t_* is selected such that the loss function *L*(˙) is minimized. For this work we have used a gradient-boosting tree classifier ensemble implementation from the “catboost” v0.10.3 R library [70]. Likewise, feature importance analysis made use of relevant implementations available as part of this library. Specifically, we have used importance analysis to evaluate overall utilization of different features within our model and SHAP value analysis to do more detailed analysis of feature contributions to classification of specific instances (safe drugs vs. idiosyncratically toxic subset identified from literature). Classifier hyper-parameters and parameters of the network feature design were tuned on the test set using grid search, then optimal configuration was validated on the hold-out set and used to train the final model. Primary evaluation of performance was done on the basis of area under the receiver-operator curve (ROC-AUC), computed using “PRROC” R package with ‘toxic’ class instances set as foreground class.

### Validation using side effects data

Side effects information was downloaded from the OFFSIDES database [39]. This drug annotation was combined with the STITCH database [58] in order to map them to protein association network. As opposed to ChEMBL, which was used to construct our training data, STITCH also includes speculative protein binding annotation. Therefore, to make these datasets comparable, only high-confidence (score > 800) annotations and proteins also present in the protein-protein association network were retained. Drugs were filtered to remove those present in either training or hold-out subsets, which resulted in 339 compounds remaining. From the set of available annotations for those drugs, we have selected all major toxicity-associated side effects with at least 10 occurrences, which resulted in fourteen categories. These side effects included most categories commonly linked to drug withdrawals [4], like cardiotoxicity, hepatotoxicity and nephropathy. Predicted scores of drugs in those categories were compared to remaining subset which did not have any of these annotations using Wilcoxon signed ranks test.

### Model interpretation and annotation of druggable proteome

The overall contribution of individual features to the model was quantified with feature importance metric and possible relationships between individual features with interaction strength metric using implementations included in the “catboost” library. In order to extract the overall map of toxicity risk from the model, we have used a druggable genome dataset from [40]. The reasoning behind this choice was that these drug-binding proteins are both most relevant and most likely to be consistent with the data used for training. For each protein, a simulated parenteral drug instance was generated using real DSD distance to the reference points and GO functional annotation of that protein, with remaining features set to missing. To explore possible patterns, distances of these proteins to twelve reference points were mapped onto a 2-D space using the t-SNE algorithm with default settings. To further profile areas of highest predicted toxicity, the top 10% of proteins by score were analyzed as a separate set. Specifically, modules in this subset were identified by fitting a Gaussian finite mixture model using the expectation-maximization algorithm. This analysis was done using an implementation from “pvclust” R package [71]. Then, functional enrichment test was done for each identified module using Fisher’s Exact Test followed by Benjamini-Hochberg False Discovery Rate correction.

An implementation of the method and supporting data have been made available in a public GitHub repository with the following URL: https://github.com/artem-lysenko/TargeTox. All other data used in this study was acquired from relevant public resources as identified in the Methods section.

## Acknowledgements

This work was supported by Core Research for Evolutional Science and Technology (CREST) Grant from the Japan Science and Technology Agency (Grant Number: JPMJCR1412), and Japan Society for the Promotion of Science (JSPS) KAKENHI (Grant Numbers: 18K18156, 17H06307 and 17H06299).

## Author Contributions

A Lysenko: conceptualization, methodology development, implementation, analysis, validation, preparation of figures and original manuscript, review and editing.

A Sharma: manuscript preparation, methodology development.

KA Boroevich: data curation, analysis, manuscript review and editing.

T Tsunoda: funding acquisition, conceptualization, investigation, supervision, project administration, manuscript review and editing.

## Conflict of Interest Statement

The authors declare no competing interests.

## Supplementary figure legends

**Figure S1 Evaluation of PrOCTOR and wQED performance on withdrawn from market subset**

Receiver-operator characteristic curve was computed from scores returned by those methods for safe and “withdrawn from market” subsets. As neither of these tools were originally designed to be used for such instances, this should not be interpreted as representative of their performance in intended context, but does suggest a potential important “blind spot” of current methods.

**Figure S2 Evaluation results for different baselines and method design variants on the training set**

Receiver-operator characteristic curve was computed on the training set (n=719) using 5-fold cross-validation. **(A)** final model that used diffusion state distance and 12 reference points **(B)** Mashup-based distance variant **(C)** all possible 16,610 reference points with diffusion state distance **(D)** distance to single medoid of all drug-binding proteins variant **(E)** Shortest path distance variant **(F)** discretized shortest path variant.

**Figure S3 Evaluation of ability to distinguish idiosyncratically toxic drugs from safe subset**

**(A)** TargeTox method scores computed using leave-one-out cross-validation **(B)** PrOCTOR scores as returned by the trained model released by the authors **(C)** weighted QED scores. In all cases exactly the same dataset was used (n = 696 in safe category and n =38 in toxic category) and in all cases significance was computed using Wilcoxon signed ranks test.

## Table Titles

**Table 1 Comparative Shapley additive explanation (SHAP) analysis of model feature importance.**

**Table 2 Average scores of OFFSIDES toxicity categories compared to 120 drugs without such annotations.**

Averdage distances between drug targets of each multi-target drug versus random samples of the same size

